# The degree of subgenome expression bias in *B. napus* changes between cultivars, tissues and across time

**DOI:** 10.64898/2026.06.01.728460

**Authors:** Hugh Woolfenden, Rachel Wells, Richard J. Morris

**Affiliations:** Computational and Systems Biology, John Innes Centre, Norwich Research Park, Norwich NR4 7UH, UK; Crop Genetics, John Innes Centre, Norwich Research Park, Norwich NR4 7UH, UK

**Keywords:** Homoeologue Expression Bias, Genome dominance, *Brassica napus*, Oilseed rape, Polyploid, Transcriptomics, Time series

## Abstract

Most extant plants show evidence of past polyploidization events in their genomes. Allopolyploids arise from hybridisation, resulting in the polyploid genome comprising subgenomes from different ancestors. Subsequent adaptation to their environment or selection pressure for specific traits has led to several allopolyploids exhibiting an unequal contribution from their subgenomes to their phenotype. Given the diversity of cultivars grown for different environments, it is possible that the associated regulatory changes may have given rise to different subgenome expression biases. Likewise, different tissues and developmental stages have distinct expression profiles that may correspond more strongly to one subgenome over the other(s). Here, we investigate different metrics for quantifying the contribution of each subgenome in space (tissue) and time (development) in cultivars of *Brassica napus*. *Brassica napus* has two subgenomes, A and C, from its ancestors *Brassica rapa* (A) and *Brassica oleracea* (C). We find that the C genome has higher overall expression than the A genome, whereas the average expression per gene is higher for the A genome. Direct comparison of homoeologous pairs reveals higher expression of genes on the C genome. We find that the degree of expression bias can change between cultivars, tissues and across time with bias quantification being strongly dependent on the metric. These findings help explain contradictory reports on expression bias and genome dominance.

**Significance statement:** We demonstrate how different metrics of expression bias between subgenomes in polyploids can lead to conflicting inferences. We show that expression can be viewed as either A or C-biased, yet the differences are small, calling into question the relevance of subgenome dominance and expression bias in *B. napus*.

## Introduction

Polyploidy is common across the tree of life, arising from whole genome duplication and/or hybridisation events (Morris *et al*., 2024). Whole genome duplication gives rise to autopolyploidy, i.e. multiple copies of the same genome, whereas allopolyploidy refers to the result of a hybridisation event between two different species and the consequent merger of the parental genomes, now termed subgenomes of the new species (Glover *et al*., 2016). Many important crops are allopolyploids, including several species of *Brassica* (oilseed rape, brown mustard), *Gossypium* (cotton), *Triticum* (wheat), and *Fragaria* (strawberry).

Studies using resynthesised polyploids (Rapp *et al*., 2009; Wu *et al*., 2018; Li *et al*., 2020; Bird *et al*., 2021) have shown that when different parental genomes come together they undergo a phase of rapid regulatory changes in which the subgenomes adapt to their new configuration (Alger and Edger, 2020). This dramatic adjustment has been described as a “transcriptome shock” (Hegarty *et al*., 2006). The first consequences of polyploidisation are changes in expression. Over longer timescales, accumulated mutations can result in the loss of genes from one or more of the subgenomes, termed fractionation, with potentially even whole chromosomes being lost (Xiong *et al*., 2011; Sankoff and Zheng, 2012). The genetic redundancy created by polyploidisation leads to genes that have been duplicated from an ancestral single gene copy being under less selection pressure, allowing for sub- and neo-functionalisation (Conant and Wolfe, 2008). These processes are thought to be major drivers for innovation and adaptation to extreme changes in the environment (Sobhanian *et al*., 2026). Likewise, selective breeding exploits the genetic redundancy of polyploids, resulting in changes in regulation that are tailored towards specific traits and climatic conditions.

As a consequence of regulatory adjustments, genes from one subgenome may become more highly expressed than the other(s) and genes inherited from the progenitors may not be lost equally across the subgenomes as the species evolves. These cases have been described as one subgenome being more “dominant” (Alger and Edger, 2020). Over evolutionary time, polyploids typically revert back to a diploid state (diploidisation) (Conant *et al*., 2014; Bird *et al*., 2018), often leaving ancestral remnants of polyploidy that can still be detected (Jiao *et al*., 2011).

Due to its agronomic importance and interesting history of polyploidisation events, the allotetraploid *Brassica napus* (oilseed rape, OSR) has been the focus of several studies on genome dominance. *Brassica napus* is the result of a hybridisation event of its progenitor species, *B. oleracea* and *B. rapa*, about 7,500 years ago (Chalhoub *et al*., 2014). Using a resynthesised line, Wu *et al*. (2018) examined homoeologue expression in leaves and found a bias to the A subgenome. Later, expanding the range of tissues to stems, leaves, flowers and siliques, a mixed picture emerged with mostly the A subgenome being dominant but in some cases the C subgenome, notably in the leaves (Li *et al*., 2020). Possible explanations for the discrepancy were discussed and included the leaf age at the time of sampling and the sampled variety (Li *et al*., 2020). In a separate study in younger leaves, the C subgenome was found to be dominant over multiple generations (Bird *et al*., 2021). The bias to the C subgenome was also found in leaf tissue when the plants were subjected to *Sclerotinia* infection (Jong and Adams, 2023). Analysis of transcript expression in individual tissues of developing seeds has also highlighted conflicting biases, with A subgenome bias inferred in one experiment (Khan *et al*., 2022), and later, bias to the C subgenome (Ziegler *et al*., 2025). This is perhaps not surprising, as gene expression is highly dynamic and tissue-specific, meaning that measures of subgenome bias based on expression may vary within a single plant. Indeed, at specific time-points the dominant subgenome may in fact be more lowly expressed (Alger and Edger, 2020), thus highlighting the importance of examining trends between tissues and across time.

Here, we quantify subgenome expression bias for a set of cultivars of *B. napus* over time and in different tissues. We investigate subgenome bias using gene expression in apex and leaf tissues across development in three *B. napus* cultivars, with each cultivar representative of a specific seasonal type. If selective breeding tailored towards climatic conditions results in changes in regulation, we might expect different cultivars to have differing degrees of adjustment in each subgenome. For instance, a rapid-cycling cultivar may draw more on the machinery from the A subgenome (Calderwood *et al*., 2021). Overall, we find that total gene expression has a larger contribution from the C subgenome. However, in terms of mean expression per gene, the A subgenome is dominant. The number of genes expressed from C exceeded the number from A, but not in proportion to the actual number of annotated genes on each subgenome, with A expressing a higher ratio of genes than C. These observations were consistent between tissues, across time and in all three genotypes. When we consider only the homoeologue gene pairs, we find an expression bias towards the C subgenome across tissues, time and genotypes.

Following previous investigations (Ramírez-González *et al*., 2018; Bird *et al*., 2021; Jong and Adams, 2023; Glombik *et al*., 2025), we examined the distributions of relative expression levels for the homoeologue gene pairs. We find that the choice of threshold used to define a homoeologue gene pair as biased can lead to inconsistent inferences, with significant bias switching between subgenomes. We found that bias amongst the homoeologue gene pairs could be transient. These temporal aspects as well as the variation between replicates, highlight the difficulties inherent in quantifying expression bias.

## Results

### Summary of a *B. napus* developmental time-course dataset

To investigate subgenome bias in *B. napus* we exploited an available RNA-seq time-course dataset (Woolfenden *et al*., 2026) from apex and leaf tissue in three cultivars (Figure 1). Both apex and leaf samples were collected across the developmental life cycle, starting from young plants (day 14 after sowing) to the time-point where first buds became visible (BBCH51). The cultivars in this dataset include a spring type (Stellar DH), a semi-winter (Zhongshuang 11) and a biennial kale (Ragged Jack). The semi-winter and biennial kale cultivars both require a period of cold (of different lengths) to induce vernalisation. The mean number of raw reads across all genotypes and tissues was 28.9 M reads, with minimum and maximum numbers, 24.2 M and 38.5 M reads, respectively. The reads were aligned to the Darmor-bzh version 10 genome reference (Rousseau-Gueutin *et al*., 2020) with the gene expression quantification for the uniquely mapped reads used herein. The mean alignment rate for the uniquely mapped reads was 82.5%. The documentation of the dataset (Woolfenden *et al*., 2026) provides full details regarding the growth conditions, sample collection, sequencing, and gene expression quantification.

**Figure 1.**
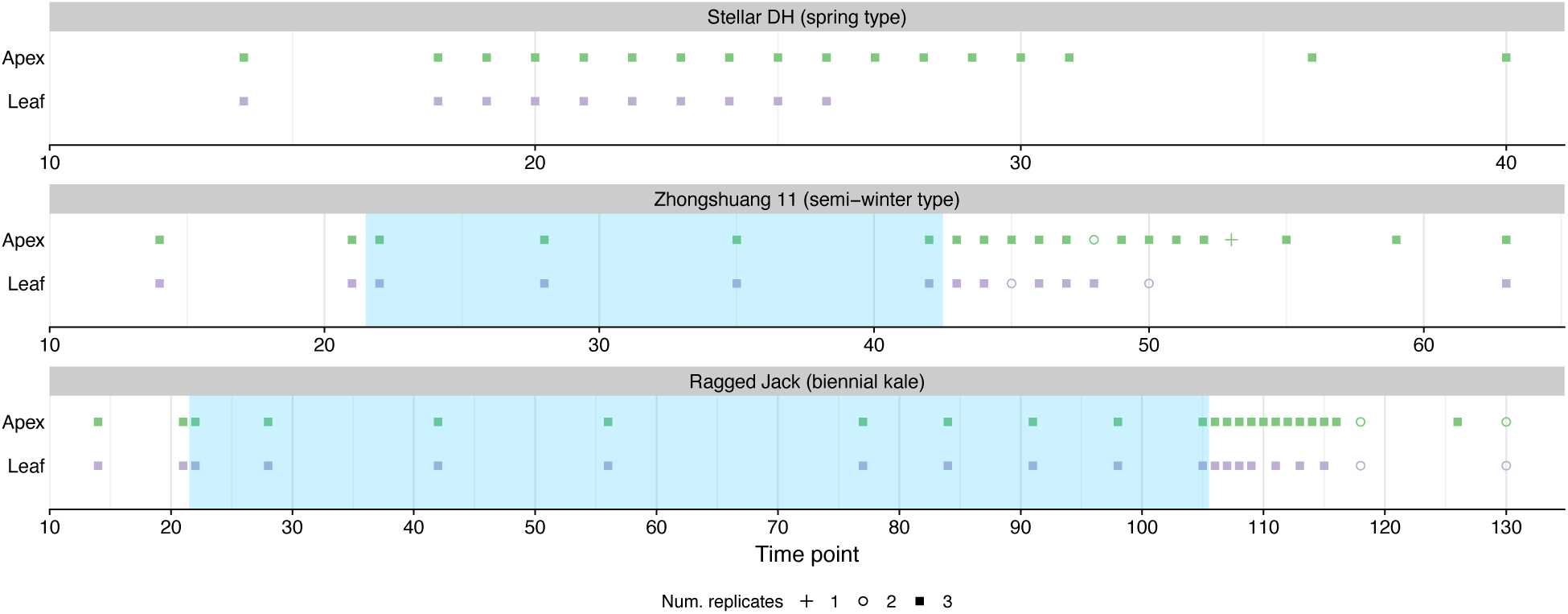
Sample time-points for RNA-seq during development for three B. napus cultivars. Apex and leaf tissues were sampled at the indicated time-points in the spring-type Stellar DH (top panel), in the semi-winter type Zhongshuang 11 (ZS) (middle panel), and in the biennial kale, Ragged Jack (RJ) (bottom panel). The number of replicates, 1, 2 or 3, is shown by the shape of the point. The period of cold to which ZS and RJ were subjected to induce vernalisation is represented by the blue shaded region. The numerical values on the horizontal axis are the number of days after sowing.

### Total gene expression from the C subgenome is higher than from the A subgenome

We first compared the total expression from the A and C subgenomes at each time-point. We calculated the total expression from each subgenome as the sum of individual TPM values for each sample (where sample uniquely identifies the genotype/tissue/time-point/replicate combination) in all time-series (Figure S1). The ratio of the expression from A over the expression from C is shown in Figure 2. Consistent with Ziegler *et al*. (2025), we see that the total expression from the C subgenome exceeds that of the A subgenome (A/C ratio less than 1). However, the difference in expression between subgenomes is not large (less than 10%).

**Figure 2.**
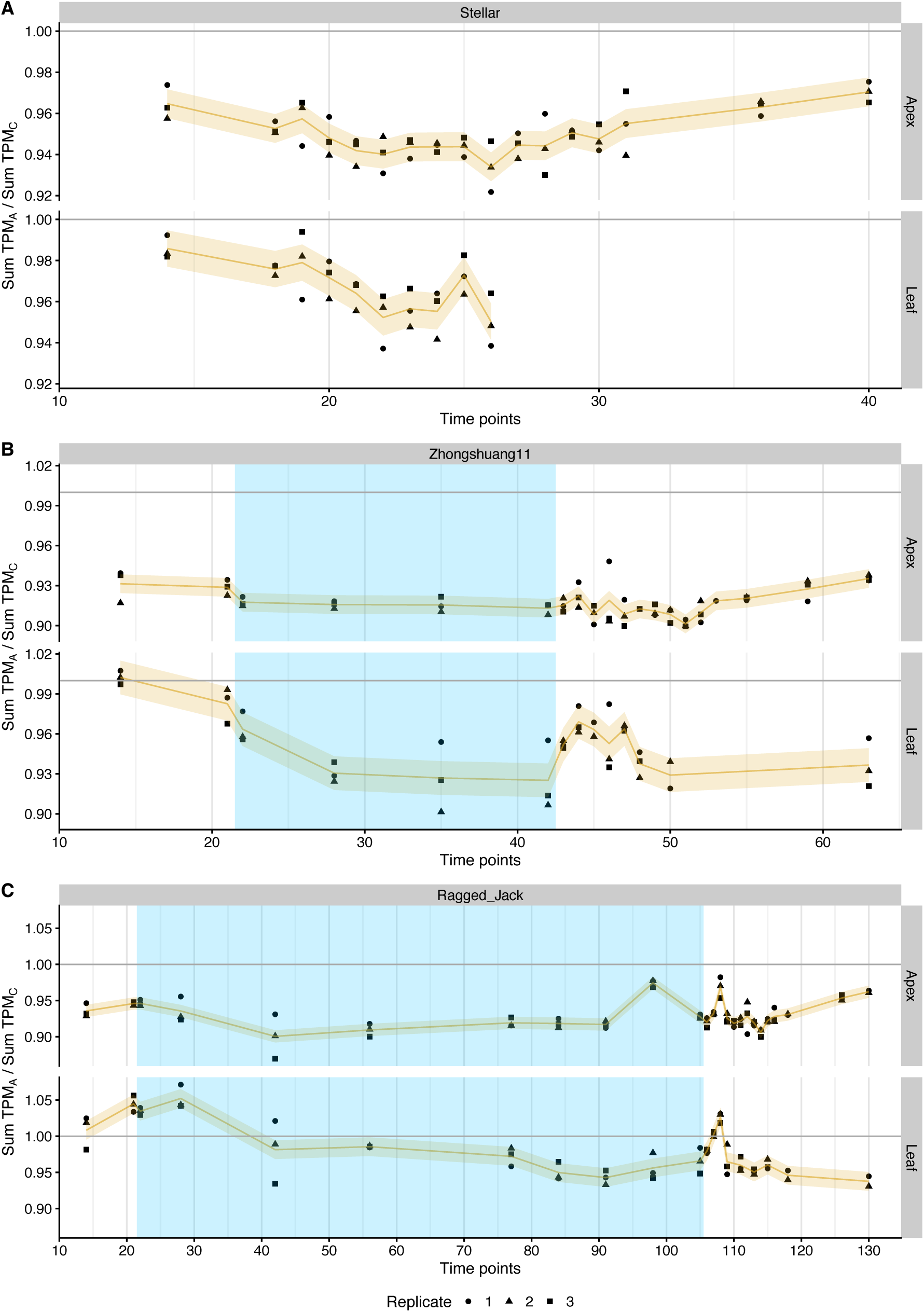
Total gene expression is biased to the C subgenome. The ratio of the sum of the gene expression values (TPM) for each subgenome across time, in both tissues, and in genotypes, Stellar (A), Zhongshuang 11 (B), and Ragged Jack (C). The period of cold treatment is depicted by the blue shaded regions. Each point represents a sample with the shape of the point indicating the replicate. The trend is shown with a thin orange line and the estimated standard deviation per time series by a light orange band (see Methods).

We next tested for tissue specificity. In the apex of Stellar, we found that expression from the C subgenome was on average 5.3% higher than from A. We denote this as 5.3% C>A. The range went from 2.5% C>A to 8.5% C>A, i.e. expression from C was higher in all samples. In Stellar leaf tissue, subgenome C was also consistently more highly expressed but less so than in the apex, 3.5% C>A (0.6% – 6.7% C>A). This trend was consistent across cultivars. In the apex for Zhongshuang 11 we found 9.1% C>A (5.5% – 11.2% C>A), whereas leaf tissue had 5.0% C>A (0.7% A>C – 10.9% C>A). In the apex of Ragged Jack the bias was 7.6% C>A (1.8% – 15.0% C>A) and 1.9% C>A (7.1% A>C – 7.4% C>A) in leaf, with some stages of development exhibiting higher expression from the A subgenome. Overall, we found that the value of C>A expression bias was significantly higher in the apex than the leaf (p-value<4e-6, Welch’s t-test) (Figure S2).

Furthermore, we found that the differences in A/C expression between cultivars were significant (p-value<2e-5 for apex and p-value<0.05 for leaf, Games-Howell pairwise test with Holm post-hoc correction), with Zhongshuang 11 showing the most pronounced subgenome expression differences.

In summary, total expression from the C subgenome was on average higher than from the A subgenome in the two tested tissue types of three cultivars of *B. napus*. The A/C expression ratios were significantly different between tissues and cultivars. However, these differences are small and there is variation within replicates and between time-points.

### The C subgenome expresses more genes, however, mean expression per gene is higher for the A subgenome

The higher expression from the C subgenome may be a consequence of different numbers of genes between subgenomes. There are 47,193 (44.4%) and 59,692 (55.6%) annotated genes on the A and C subgenomes (Darmor-bzh, version 10), respectively. The difference in the number of expressed genes (TPM>1) is consistent between subgenomes across tissues and cultivars (Figure 3, Figure S3). The A/C ratio is less than 1 at every time-point in all three genotypes (Figure 3, Figure S3). This agrees with previous findings (Ziegler *et al*., 2025).

**Figure 3.**
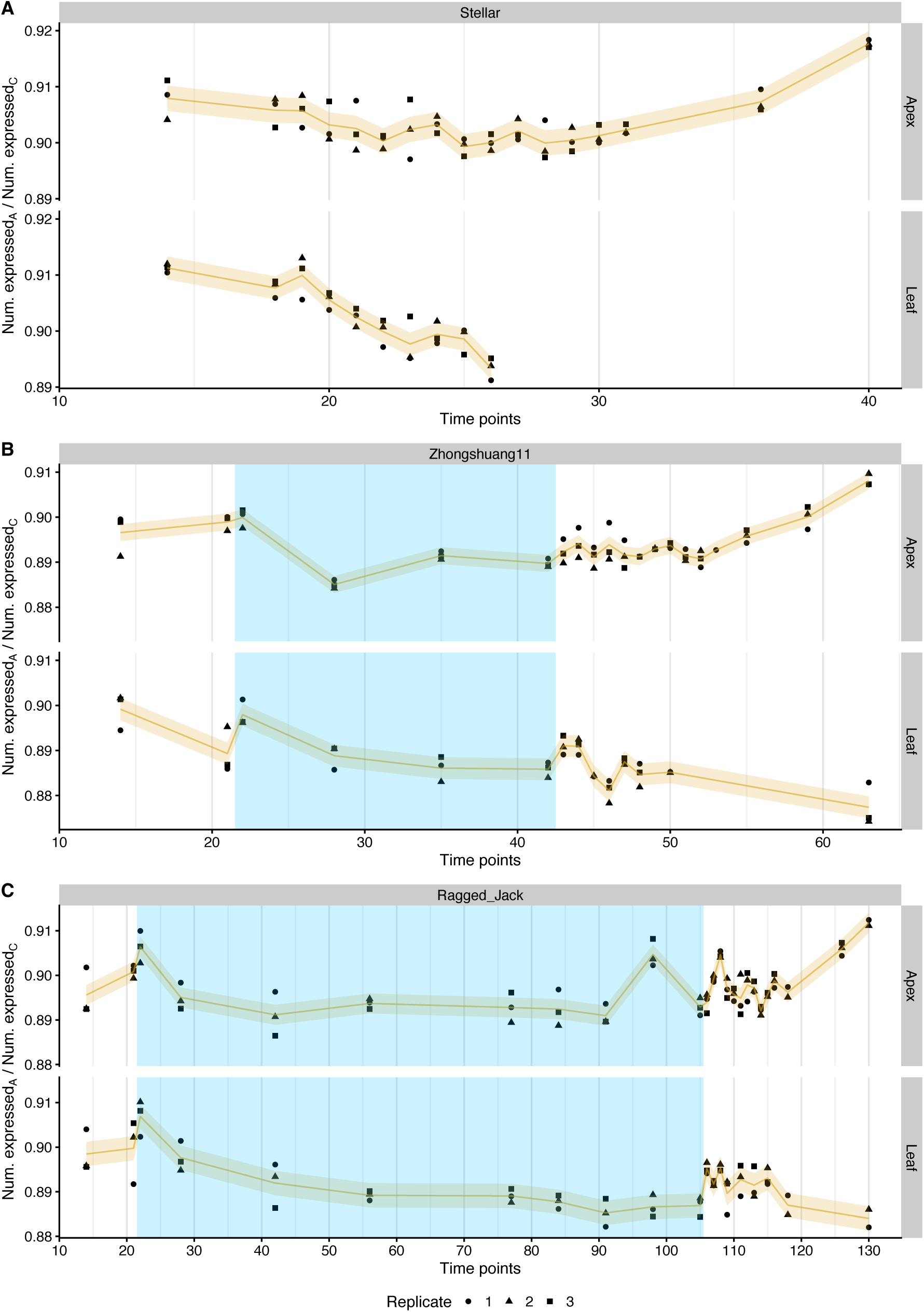
The C subgenome consistently expresses more genes than the A subgenome. The ratio of the number of expressed genes on the A and C subgenomes across time, in both tissues, and in genotypes, Stellar (A), Zhongshuang11 (B) and Ragged Jack (C). Based on these numbers we find bias in Stellar of 10.7% C>A (8.9% -11.5% C>A) in the apex and 10.8% C>A (9.5% -12.2% C>A) in the leaf, for Zhongshuang 11 we find 11.8% C>A (9.9% -13.1% C>A) in the apex and 12.6% C>A (10.9% – 14.4% C>A) in the leaf, and for Ragged Jack the apex had 11.4% C>A (9.6% -12.8% C>A) and the leaf 12.1% C>A (9.9% – 13.4% C>A). The period of cold treatment is depicted by the blue shaded regions. Each point represents a sample with the shape of the point indicating the replicate. The trend is shown with a thin orange line and the estimated standard deviation per time series by a light orange band (see Methods). A gene was considered to be expressed if its TPM was above 1.

The A/C ratios show variability between tissue and genotype (Figure S4). The ratio of the number of expressed genes between apex and leaf is significantly different (p-value<3e-6, Welch’s t-test) for Zhongshuang 11 and Ragged Jack, and between the three genotypes in each tissue (p-value<4e-3, Games-Howell pairwise test with Holm post-hoc correction).

Based on the number of annotated genes, we expected the number of expressed genes to be 26% C>A (=59,692/47,193 – 1) but our observations above deviated from this. We therefore checked to see if the numbers of expressed genes on the two subgenomes diverged from this significantly. We found the number of expressed genes was significantly higher than expected for A and lower than expected for C (p-value<1e-24, 𝜒^!^ test). Given transposons are more numerous on the C subgenome (Chalhoub *et al*., 2014) this could be an effect of transposon-related expression suppression.

We computed the average TPM per expressed gene for each subgenome by dividing the total expression value by the number of expressed genes (Figure S5). The mean TPM per expressed gene was consistently greater for subgenome A than C (Figure 4). In the apex, the mean expression from subgenome A was 5.2%, 2.6% and 3.7% higher than from C in Stellar, Zhongshuang 11 and Ragged Jack, respectively. These values are significantly lower (p-value<2e-9, Welch’s t-test) than in the leaf, where we found 7.2%, 7.4%, and 10.2% for the cultivars Stellar, Zhongshuang 11 and Ragged Jack, respectively.

**Figure 4.**
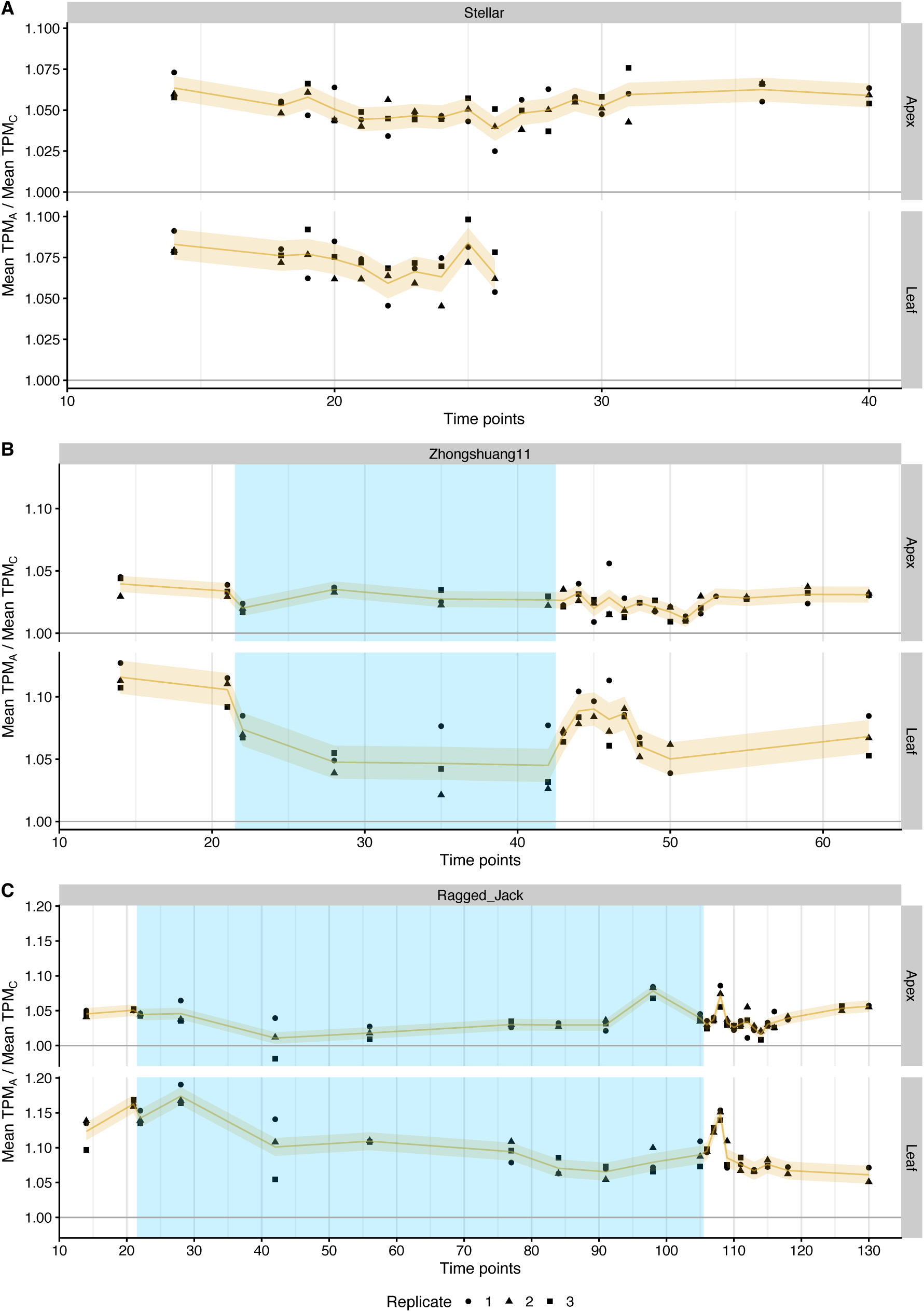
The mean expression per gene is higher for the A subgenome. The ratio of the mean of the gene expression for expressed genes by subgenome across time, in both tissues, and in genotypes, Stellar (A), Zhongshuang 11 (B), and Ragged Jack (C). The period of cold treatment is depicted by the blue shaded regions. Each point represents a sample with the shape of the point indicating the replicate. The trend is shown with a thin orange line and the estimated standard deviation per time series by a light orange band (see Methods). A gene was considered to be expressed if its TPM was above 1.

Apart from Stellar and Zhongshuang 11 in leaf, the cultivars exhibited a significantly different (p-value<2e-4, Games-Howell pairwise test with Holm post-hoc correction) in mean expression between subgenomes within each tissue (Figure S6).

In summary, the number of expressed genes from the C subgenome is higher, while the mean TPM per gene is higher in the A subgenome. We find significant differences between tissues and between cultivars. However, these differences are typically well within 10%.

### Homoeologue expression is higher from the C subgenome

Following our observation that total expression is higher from the C subgenome yet mean expression per gene is higher for the A subgenome, we sought to directly compare genes common to both subgenomes using homoeologue gene pairs (HGPs).

To see how expression varied between two genes in a HGP we first examined a single sample for each tissue (Stellar replicate 1 at day 14). We computed the relative expression (RE = TPM_A_/(TPM_A_ + TPM_C_)) (see Methods) per HGP, with the minimal requirement that at least one gene was expressed (TPM>1), Figure 5A. The bulk of the distribution is centred around 0.5 (equal TPM values). There are, however, large peaks at the extremities (extreme RE), with zero associated with extreme C bias and one with extreme A bias. HGPs with RE at (or close to) these extreme values could arise when only one gene is responsible for function with the other gene effectively silenced.

**Figure 5.**
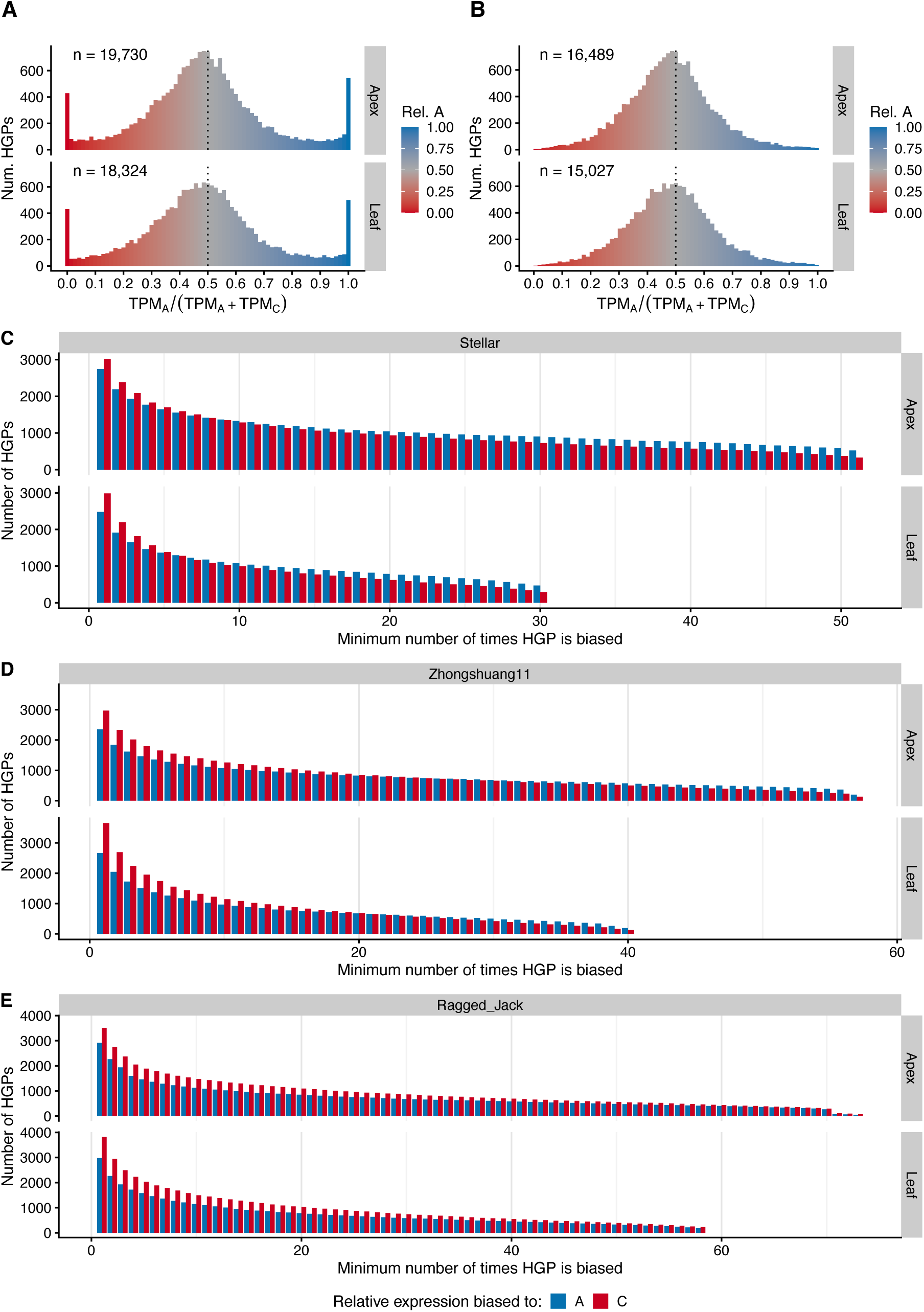
Very few HGPs exhibit consistent extreme bias in which one gene is not expressed whilst the other is. (A) The distribution of relative expression (TPM_A_/(TPM_A_ + TPM_C_)) for replicate 1 of Stellar apex and leaf at day 14. Values close to 0 and 1 indicate high C and A bias, respectively, with 0.5 occurring for equally expressed genes in the HGP. The number of HGPs in the distribution is shown at the top-left. Only HGPs with at least one expressed gene (TPM>1) are included. (B) Same as (A) except that both genes in the HGP must have TPM values above one. (C) The number of HGPs (vertical axis) that had extreme relative expression (below 0.1 or above 0.9) in a minimum number of samples (horizontal axis) for Stellar apex and leaf. For example, if a HGP had extreme bias in one or more samples (cultivar/tissue/time-point/replicate) then it would be included in the left-most set, whereas a HGP that is biased in every sample (“core”) would be included in the right-most set. Only HGPs with at least one expressed gene (TPM>1) are included. The extreme bias is to A or C, which is indicated by the colour (A bias in blue and C bias in red). There were 392 A-biased and 221 C-biased (392 A/221 C) “core” HGPs common to apex and leaf, which formed part of 523 A/326 C in apex and 472 A/293 C in leaf. (D) Similar to (C) for Zhongshuang 11. There were 197 A/130 C “core” HGPs in apex, 190 A/122 C in leaf, and 121 A/62 C were common to both tissues. (E) Similar to (C) for Ragged Jack. There were 52 A/74 C “core” HGPs in apex, 185 A/228 C in leaf, and 26 A/46 C were common. The number of “core” HGPs common to all six time-series was 24 (7 A/17 C).

To quantify how many HGPs fell into this silenced category, characterised by extreme RE values, we counted the number of HGPs with RE below 0.1 or above 0.9 across samples. Our classification of HGP as exhibiting extreme bias is similar to previous investigations (Bird *et al*., 2021; Jong and Adams, 2023; Ziegler *et al*., 2025) that classified HGP bias into three bias classes (A-biased, C-biased, or not biased). We found that around 6,000 HGPs had extreme RE in one or more samples (in each of the time-series) but that this expression bias was not consistent over time (Figure 5C-E). We found 392 HGPs with expression only from the A subgenome and 221 only from C in Stellar.

To investigate whether specific biological processes, molecular functions, or cellular components had undergone this extreme expression bias between subgenomes, we performed a GO-term analysis (see Methods). In the sets of core extreme HGPs, we identified no biological process terms, two significant molecular function and four cellular components that were enriched, Table S2. The identified terms do not support specific pathways or processes being selectively chosen from either subgenome. We nevertheless repeated the same analysis for the other cultivars to see whether differences could be identified that may link to their life-histories. The size of the core sets of extreme RE were smaller for the other two cultivars. For Zhongshuang 11, we identified 121 HGPs exclusively expressed from A and 62 from C, whereas only 26 in A and 46 in C in Ragged Jack. For Zhongshuang 11, GO-term analysis only found two significantly enriched terms, Table S2. There was a lack of consistency between the terms for Stellar and Zhongshuang 11, with “structural molecule activity” being the only common term. There was no significant enrichment for Ragged Jack. The GO-term analysis found significant enrichment but no pattern to support the selection of defined pathways or processes, Table S2. In fact, the relative sizes of these core extreme HGP sets largely reflects the number of samples in each timeseries.

As other approaches have used a range of different thresholds for quantifying HGP expression bias, we repeated these analyses and investigated the impact of threshold choices. We found that a different fold-change cut-off (FCC) could result in samples that were significantly biased to C for a low FCC either losing the significance for a higher FCC, or even reversing the result so that some samples were significantly biased to A. We found that this reversal could also happen when we changed either the initial expression filter or the bias quantification method. This analysis highlights the possibility of different HGP bias classification methods producing different overall bias inferences. See Figure S7 and S8 for further detail.

In short, the extremely biased HGPs exhibit little consistency over a time-course and between cultivars. This indicates that the cases of extreme bias in HGPs are transient, and the sets observed for one sample could be highly dissimilar to the sets observed in another sample. We next extended the analysis to all HGPs where both genes were expressed (TPM > 1) and focussed on the non-extreme RE cases, Figure 5B. To quantify the overall subgenome bias, we computed the (geometric) mean over the ratios of A/C expression for all HGPs. The mean A/C ratios were all less than one (Figure 6), supporting an overall bias to the C subgenome. On average, genes in HGPs from the C subgenome were expressed 10.3% higher than genes from the A subgenome. Consistent with our previous observations using different metrics, we found a tissue-specific difference in expression bias across all cultivars, with the apex showing higher expression from C to A in HGPs than the leaf (p-value<2e-3, Welch’s t-test) (Figure S9). We also find that the different cultivars had a significantly different bias with Zhongshuang 11 being more C-biased than Stellar, which was more C-biased than Ragged Jack, in both tissues (p-value<4e-3, Games-Howell pairwise test with Holm post-hoc correction) (Figure S9).

**Figure 6.**
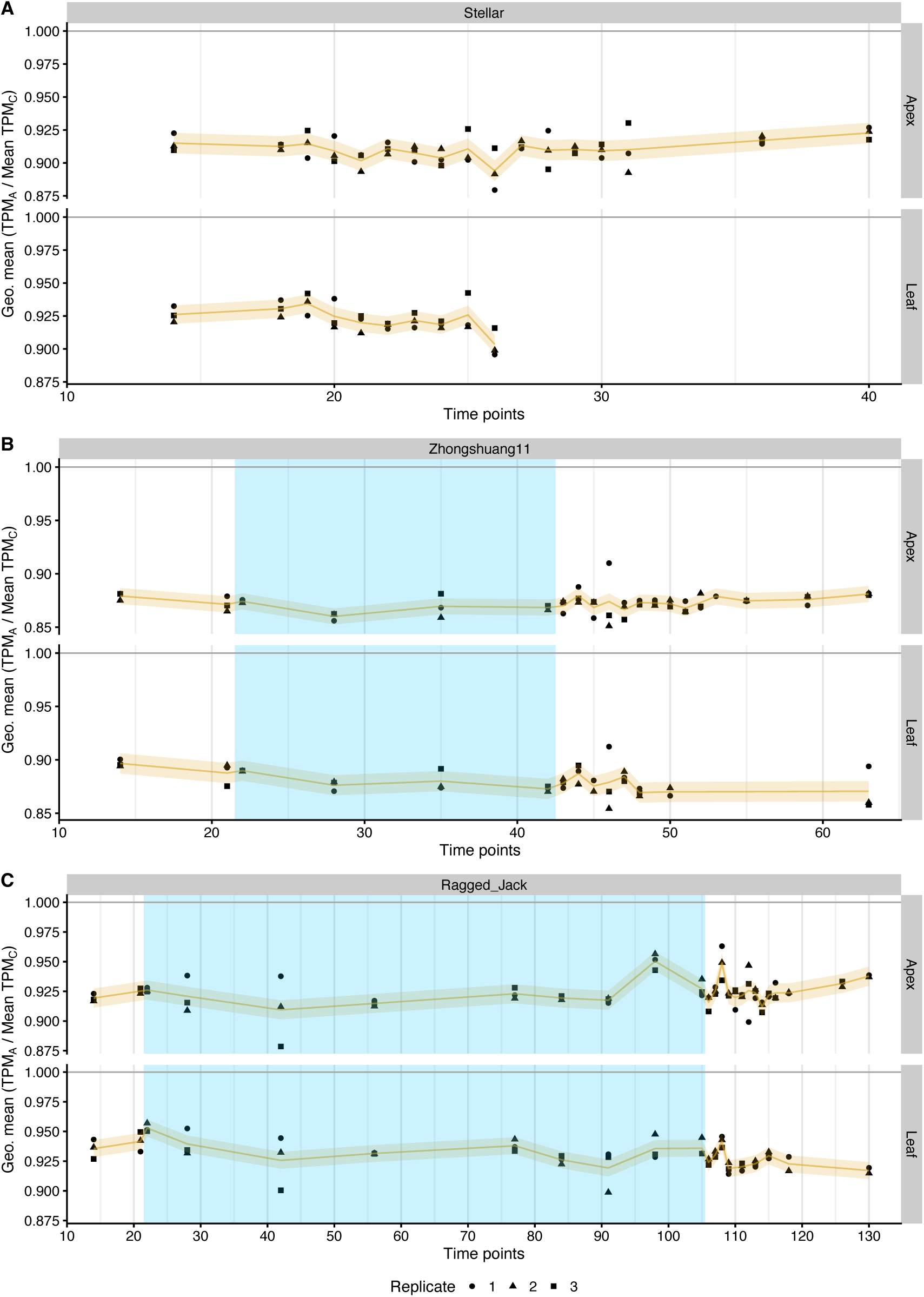
The expression of homoeologue gene pairs is higher from the C genome. Each point shows the geometric mean of the expression ratio for the A and C gene in each expressed homoeologue gene pair (HGP) across time, in both tissues, and in genotypes, Stellar (A), Zhongshuang 11 (B), and Ragged Jack (C). The mean biases for Stellar, Zhongshuang 11 and Ragged Jack were 9.9% C>A (7.5% – 13.7% C>A), 14.7% C>A (9.9% – 17.5% C>A) and 8.2% C>A (3.8% – 13.8% C>A) in the apex, respectively, and 8.4% C>A (6.1% – 11.6% C>A), 13.6% C>A (9.6% – 17.1% C>A), and 7.5% C>A (4.5% – 11.3% C>A) in the leaf, respectively. Different replicates are represented by shapes. The period of cold treatment is depicted by the blue shaded regions. The trend is shown with a thin orange line and the estimated standard deviation per time series by a light orange band (see Methods). A HGP is considered to be expressed if both genes in the HGP have a TPM above 1.

In summary, we found that even though HGPs could exhibit extreme bias (i.e., expressed vs non-expressed) only a few HGPs maintained this across a time-course with a small proportion (0.1%) common across time, tissue and genotype. When considering all expressed HGPs, we find that homoeologous genes from the C subgenome are ∼10% more highly expressed than corresponding genes from the A subgenome. Furthermore, the subgenome expression bias of HGPs is statistically significant different between tissues and between genotypes. However, we find no evidence for selection of *B. napus* cultivars influencing biological function, or preferentially silencing individual gene copies within a HGP.

## Discussion

The aftermath of a polyploidisation event is, over time, expected to give rise to adaptations in the form of regulatory changes, gene loss, and sub- and neo-functionalisation. These adaptations are likely to be different for plants grown in different environments or that were selected for specific traits. The resulting changes in regulation may lead to expression differences between subgenomes and homoeologues, known as subgenome dominance.

As expression is dynamic and can differ between tissues and cultivars, this may potentially lead to variation in subgenome bias. To investigate spatiotemporal aspects of subgenome expression bias in the allotetraploid *B. napus*, we examined gene expression in apex and leaf tissues over time in three cultivars that represented different seasonal types.

*B. napus* has two subgenomes, A and C, with subgenome A being contributed from its ancestor *B. rapa* and subgenome C from *B. oleracea*. We found that the total gene expression per subgenome was slightly larger (less than 10%) from the C subgenome than from A, consistent with the observations of Ziegler *et al*., 2025 in individual tissues of the developing *B. napus* seed. This bias in total gene expression between A and C was statistically significantly different between tissues and between cultivars. As the C subgenome contains and consistently expresses more genes than the A subgenome, higher total expression from C is not unexpected. Normalising the total gene expression per subgenome by the number of expressed genes for each subgenome, however, reveals a higher mean gene expression for genes from the A subgenome, between 2.6% and 10.2% (on average) higher than C. The lower mean expression per gene from the C subgenome might be a consequence of the greater level of transposon activity. For most combinations, we found that the mean expression per gene was significantly different between tissues within each cultivar, and between cultivars for each tissue. The metrics of total expression and mean expression per gene thus result in apparent biases to different subgenomes.

Homoeologue gene pairs (HGPs), as examples of parallel genes being retained on the progenitor genomes since their diversification from their common ancestor, allow direct comparison between A and C subgenome expression. In principle, HGPs provide the opportunity to investigate extreme cases of expression bias, where one of the genes in the HGP is expressed while the other is not. However, few such cases could be identified. Most cases of no expression of a gene in an HGP were transient, resulting in HGPs that changed expression bias from A to C, and *vice versa*, between tissues and cultivars. The mean relative expression between genes in homoeologue gene pairs (HGPs), revealed a bias to the C subgenome, with genes from C expressed on average 10.3% higher than genes from A. Thus, whilst the mean expression per gene is higher for genes from the A subgenome, the HGPs exhibit a higher expression for genes from C. These seemingly conflicting expression biases suggest that genes that are not part of a HGP must have an even greater expression bias to the A subgenome to produce the average that we observe (Figure S11). This could be linked to the life histories of the progenitors and their evolution prior to the hybridisation event, or the “transcriptome shock” experienced upon polyploidisation. Resynthesised cultivars that “replay the tape” are an excellent route into studying the latter, with the inclusion of further organs, tissues and/or time-points augmenting the spatiotemporal picture of subgenome bias. Combined with cultivar-specific genome references, such data could underpin the investigation of regulatory elements that are causally implicated in gene expression (Rockenbach *et al*., 2025).

In summary, depending on which metric was used, we found evidence for subgenome expression bias to both the A and the C subgenomes of *B. napus*. All metrics revealed differences between apex and leaf tissue, and between cultivars for a particular tissue. These results highlight the importance of examining subgenome expression not only across time and within tissues (Alger and Edger, 2020) but also between cultivars. The tissue- and genotype-specific dependencies coupled with expression dynamics, introduce challenges for comparing statements on subgenome bias between studies. For instance, previous work has demonstrated a bias to the A subgenome (Wu *et al*., 2018; Li *et al*., 2020) whereas another report identified a bias to the C subgenome (Bird *et al*., 2021; Jong and Adams, 2023). Comparisons are further complicated by the use of different expression filters and thresholds. To classify an HGP as biased requires the introduction of a threshold for the difference in expression. We found that the number of biased HGPs is sensitive to the classification method and the chosen thresholds, meaning that for comparisons between previous reports, it would be necessary to first fully harmonise the methodologies used to classify HGPs and genome dominance.

## Materials and methods

### Gene expression data

The gene expression data is from an available *B. napus* RNA-seq time-course dataset (Woolfenden *et al*., 2026). The three cultivars in this dataset include a spring type (Stellar DH), a semi-winter (Zhongshuang 11) and a biennial kale (Ragged Jack). The data were aligned to the Darmor-bzh version 10 genome reference (Rousseau-Gueutin *et al*., 2020) using HISAT2 (v2.1.0) (Kim *et al*., 2019) after the reads were trimmed with Trimmomatic (v0.39) (Bolger *et al*., 2014). Uniquely mapped reads were extracted with samtools (v1.17) (Danecek *et al*., 2021) and gene expression quantification was obtained using StringTie (v2.1.1) (Pertea *et al*., 2016). The mean number of raw reads across all genotypes and tissues was 28.9 M reads, with minimum and maximum numbers, 24.2 M and 38.5 M reads, respectively. The mean alignment rate for the uniquely mapped reads was 82.5%. The documentation of the dataset provides full details regarding the growth conditions, sample collection, sequencing, and gene expression quantification.

### Time-course analysis

Quantification for the sample replicates in each time-course are shown individually. To quantify the standard deviation (SD) for the time-courses we de-trended the value for each replicate by computing the offset to the mean of the replicates at each time-point and then computed the SD of all of the resulting values. This resulted in a single value that was then used to depict the error estimate of the data across the time-course.

### Homoeologue identification

To identify homoeologue gene pairs (HGPs) in the Darmor-bzh v10 genome reference (Rousseau-Gueutin *et al*., 2020) we first split the genome reference by subgenome and ran the mmseqs2 (Steinegger and Söding, 2017) reciprocal best hit pipeline (“easy-rbh”) between the A and C subgenomes for both the nucleotide and protein sequences with parameters “-s 7.5 - c 0.6”. This resulted in 31,214 and 32,868 potential HGPs for the nucleotide and protein sequences, respectively, and a combined total of 34,713. To account for homoeologous exchange between the subgenomes we used GENESPACE v1.3.1 (Lovell *et al*., 2022) to identify syntenic hits, and used the results to filter the list of potential HGPs to obtain the 29,336 HGPs used in the analysis. The chromosome-by-chromosome HGP counts are shown in Figure S10 and listed in Table S1.

### Homoeologue gene pair (HGP) bias classification

HGP bias classification followed a three-step process. First, the set of HGPs was obtained by filtering on the expression of the two genes in the pair. Following previous investigations, we used two filtering criteria: both genes in the pair had TPM values larger than one (labelled “A>1 & C>1”); the sum of both TPM values was larger than two (labelled “A+C>2”). The sum filter is less stringent and admits all HGPs that pass the individual TPM filter, while also including HGPs with lowly expressed genes. The second step assessed the numerical value of the relative expression. We used different methods that have all been used in previous work: the logarithm of the fold change (log_2_(𝑇𝑃𝑀_*A*_/𝑇𝑃𝑀_*C*_)), labelled “log A/C”; a “pseudo-TPM” log fold-change (log_2_((𝑇𝑃𝑀_*A*_ + 1)/(𝑇𝑃𝑀_*C*_ + 1))), “pseudo-log A/C”; a relative expression value in the range zero to one (𝑇𝑃𝑀_*A*_/(𝑇𝑃𝑀_*A*_ + 𝑇𝑃𝑀_*C*_), “rel. AC”. The log-fold change approaches have been used in *B. napus* (Bird *et al*., 2021; Jong and Adams, 2023; Ziegler *et al*., 2025) and the “rel. AC” method in wheat (Ramírez-González *et al*., 2018; Glombik *et al*., 2025). Third, the relative expression is compared to a fold-change cut-off (FCC) to classify whether the HGP is biased and to which subgenome, i.e., A-biased, C-biased, or not biased. For example, values produced by the “log A/C” method that are larger than log_2_(FCC) are labelled A-biased while values lower than −log_2_(FCC) are labelled C-biased. We used the values 2, 3 and 4 for the FCC. The FCC can be transformed to a value suitable for the “rel. AC” method by the formula, rFCC=FCC/(FCC+1), where rFCC is the “rel. AC FCC”. For example, an FCC of 3 would have a rFCC of 3/4 so that HGPs with “rel. AC” above this would be labelled as A-biased, with HGPs that have a rFCC below 1/4 (= 1 − 3/4) labelled as C-biased. It should be noted that for the “A>1 & C>1” filter the methods “log A/C” and “rel. AC” are interchangeable, and so produce the same bias classes. We thus used two different filters for expression, three methods to calculate the relative expression, and three different FCC values to investigate the robustness of each one by comparing their outcomes across a time-course. For each sample we counted the number of HGPs that were biased to each subgenome and performed a statistical test (𝜒^!^-test) to check whether the number of HGPs significantly deviated from parity (see Discussion and Figure S8).

### Gene Ontology (GO) analysis

We performed GO analysis using g:Profiler (Kolberg *et al*., 2023) (version e114_eg62_p19_27110d83) with the default settings. For input, we used the *Arabidopsis thaliana* genes corresponding to the genes of interest. We performed the homology lookup from *B. napus* to Arabidopsis using the homology data downloaded from PLAZA Plants (version 5) (Bel *et al*., 2021), from where we also downloaded the Darmor-bzh v10 (Rousseau-Gueutin *et al*., 2020) genome reference. The background set of *B. napus* genes was obtained by only taking those *B. napus* genes that were expressed (TPM>1) at least once during the relevant time-course (e.g., Stellar / Apex). We then obtained the Arabidopsis gene for these genes from the homology data. The fold enrichment is not included explicitly in the g:Profiler output but can be calculated by dividing the observed frequency (intersection size/query size) by the expected frequency (GO term size/effective domain size) where the terms in parentheses are available in the g:Profiler results.

## Data statement

The raw RNA-seq data (FASTQ files) have been deposited in the European Nucleotide Archive (ENA) under accession numbers PRJEB106353 (Stellar), PRJEB106354 (Zhongshuang 11) and PRJEB106340 (Ragged Jack). Full details of the dataset are given in (Woolfenden *et al*., 2026), which also provides the expression data for each genotype.

## Acknowledgements

HW and RM acknowledge Dr. Amy Briffa for discussions regarding the estimation of the time-course standard deviation. HW, RM and RW are grateful for support from BBSRC’s Institute Strategic Programme Genes in the Environment (BB/P013511/1) and the Institute Strategic Programme Building Resilience in Crops (BB/X01102X/1). RM acknowledges support from BBSRC’s Institute Strategic Programme on Biotic Interactions underpinning Crop Productivity (BB/J004553/1) and Plant Health (BB/P012574/1). RM and RW acknowledge support from a BBSRC Strategic Longer and Larger Grant ‘BRAVO’ (BB/P003095/1). The funders had no role in the design of the study and collection, analysis, and interpretation of data.

## Authors’ contributions

RM and RW conceived the project.

HW performed the analysis and drafted the manuscript.

All authors contributed to writing the manuscript.

The authors read and approved the final manuscript.

## Ethics approval and consent to participate

Not applicable.

## Consent for publication

Not applicable.

## Supplemental

*Table S1. List of homoeologue gene pairs. List of pairs of genes where the first gene is on the A subgenome and the second is on the C subgenome. The HGPs are within Darmor-bzh version 10 (Rousseau-Gueutin et al., 2020) (see Methods)*.

*Table S2: List of significant Gene Ontology (GO) terms for the extreme “core” homoeologue gene pairs. The “core” HGPs are defined as those with relative expression below 0.1 or above 0.9 (see Methods)*.

**Figure S1.**
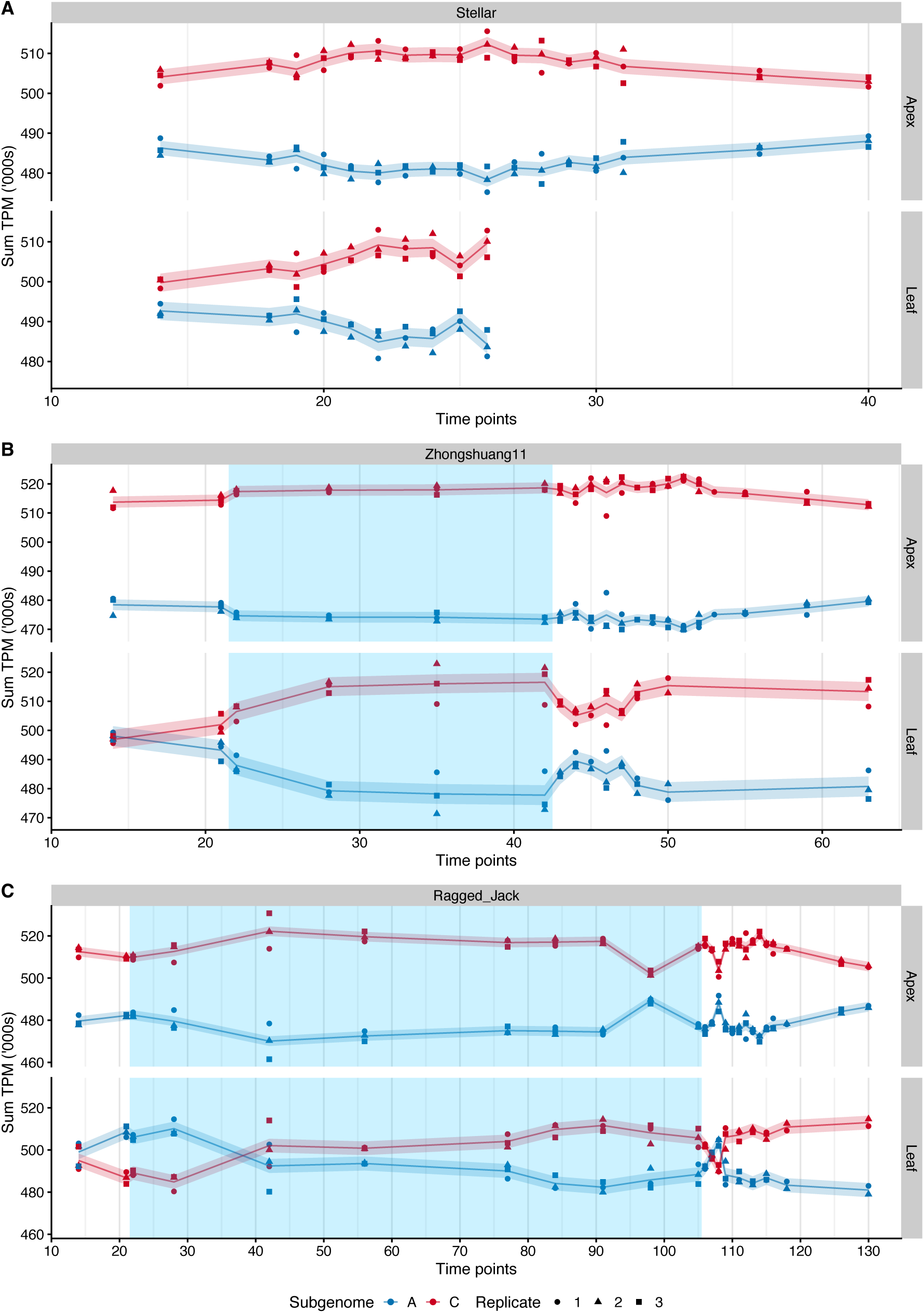
Total gene expression is biased to the C subgenome. The sum of the gene expression values (TPM) by subgenome in both tissues, and in genotypes, Stellar (A), Zhongshuang11 (B), and Ragged Jack (C). The period of cold treatment is depicted by the blue shaded regions. Each point represents a sample with the shape of the point indicating the replicate. The trend is shown with a thin line and the estimated standard deviation per time series by a shaded band (see Methods), with the colour indicating the subgenome (A: blue, C: red).

**Figure S2.**
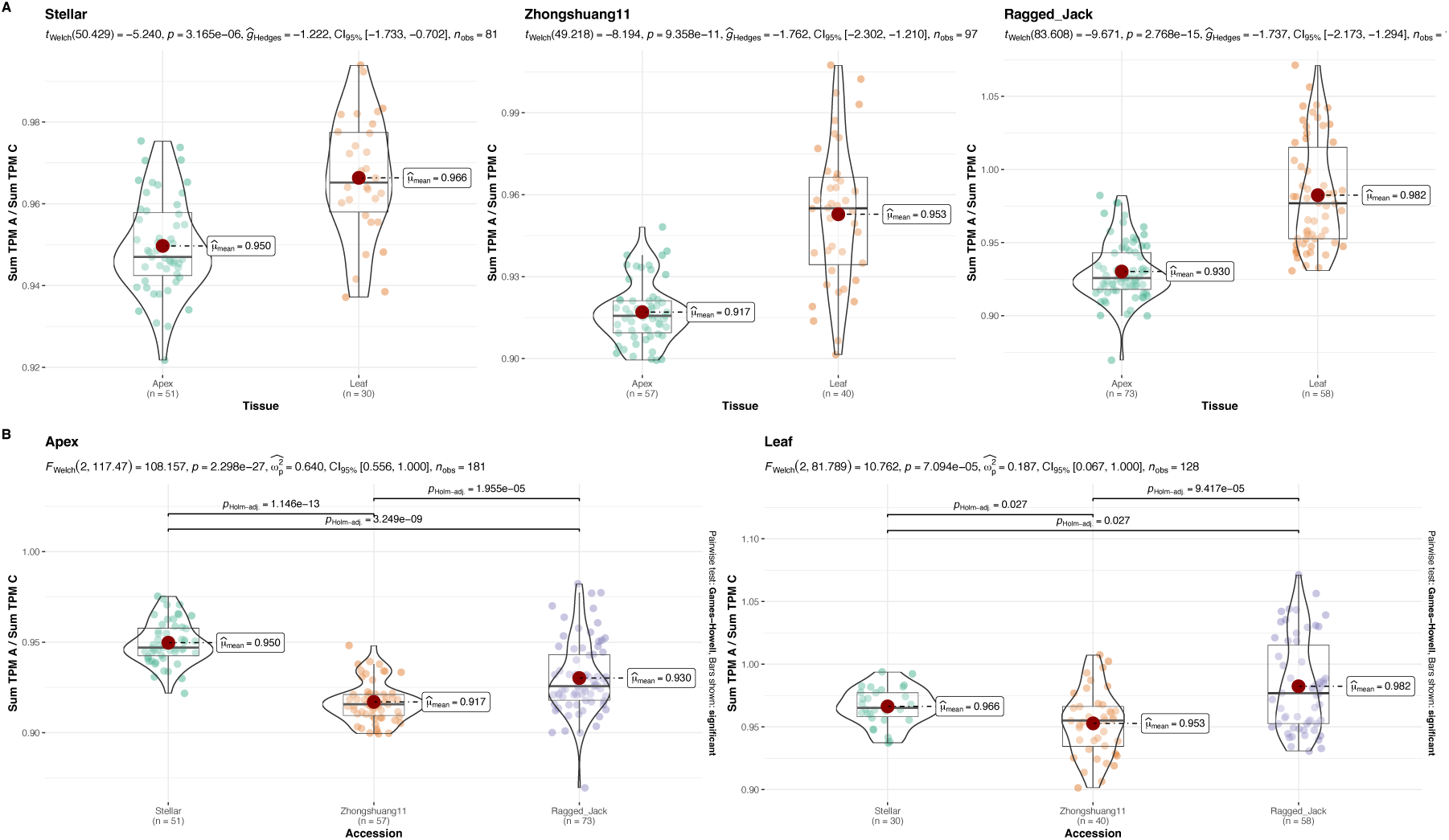
Comparison of total expression ratio between tissues and cultivars. (A) Comparison between tissues for each of the cultivars, Ragged Jack, Zhongshuang 11, and Stellar. (B) Comparison between cultivars for each of the tissues.

**Figure S3.**
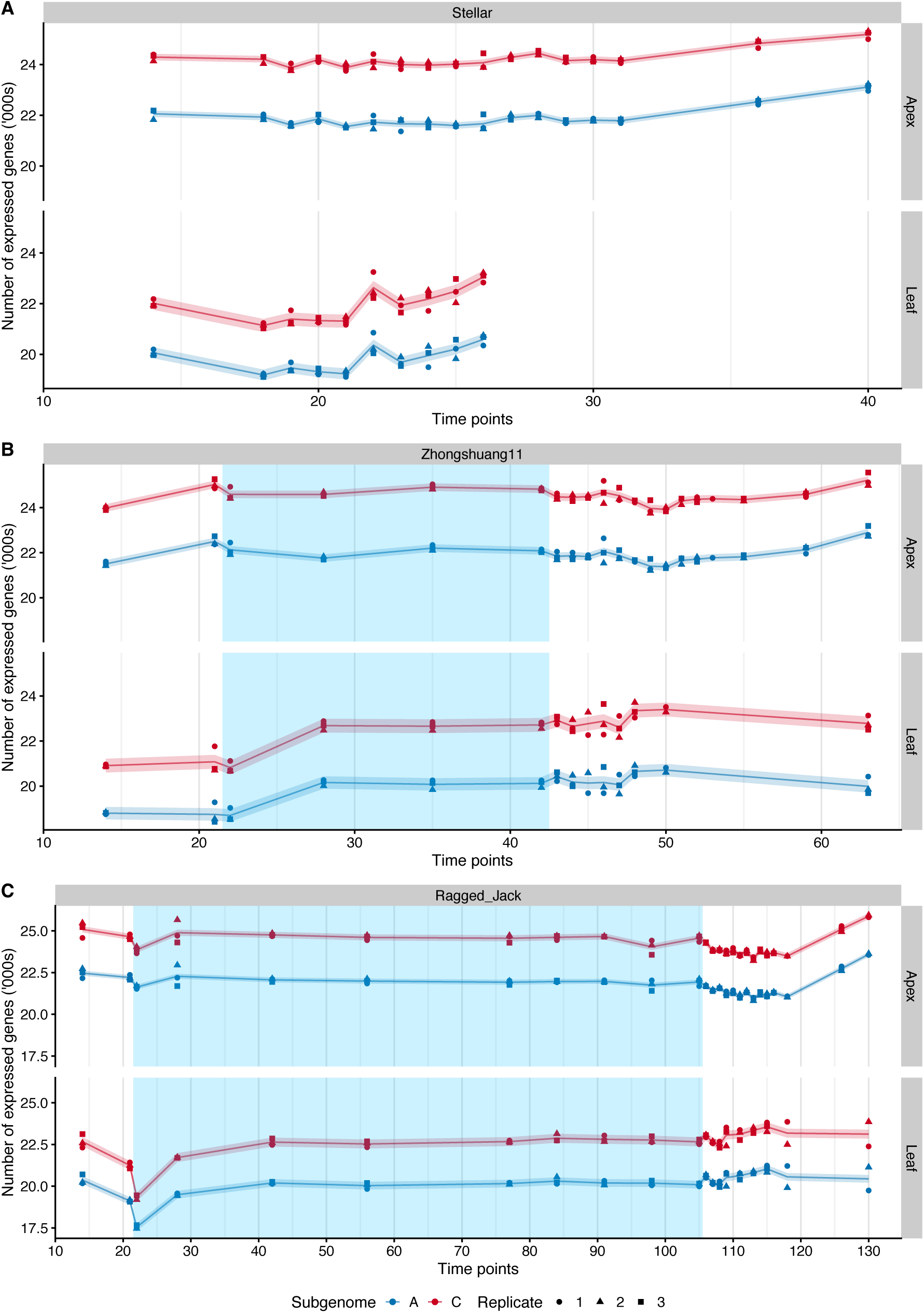
The C subgenome has consistently more expressed genes than the A subgenome. The number of expressed genes on the A and C subgenomes in Stellar (A), Zhongshuang11 (B) and Ragged Jack (C). A gene is considered to be expressed if its TPM is above 1. The period of cold treatment is depicted by the blue shaded regions. Each point represents a sample with the shape of the point indicating the replicate. The trend is shown with a thin line and the estimated standard deviation per time series by a shaded band (see Methods), with the colour indicating the subgenome (A: blue, C: red).

**Figure S4.**
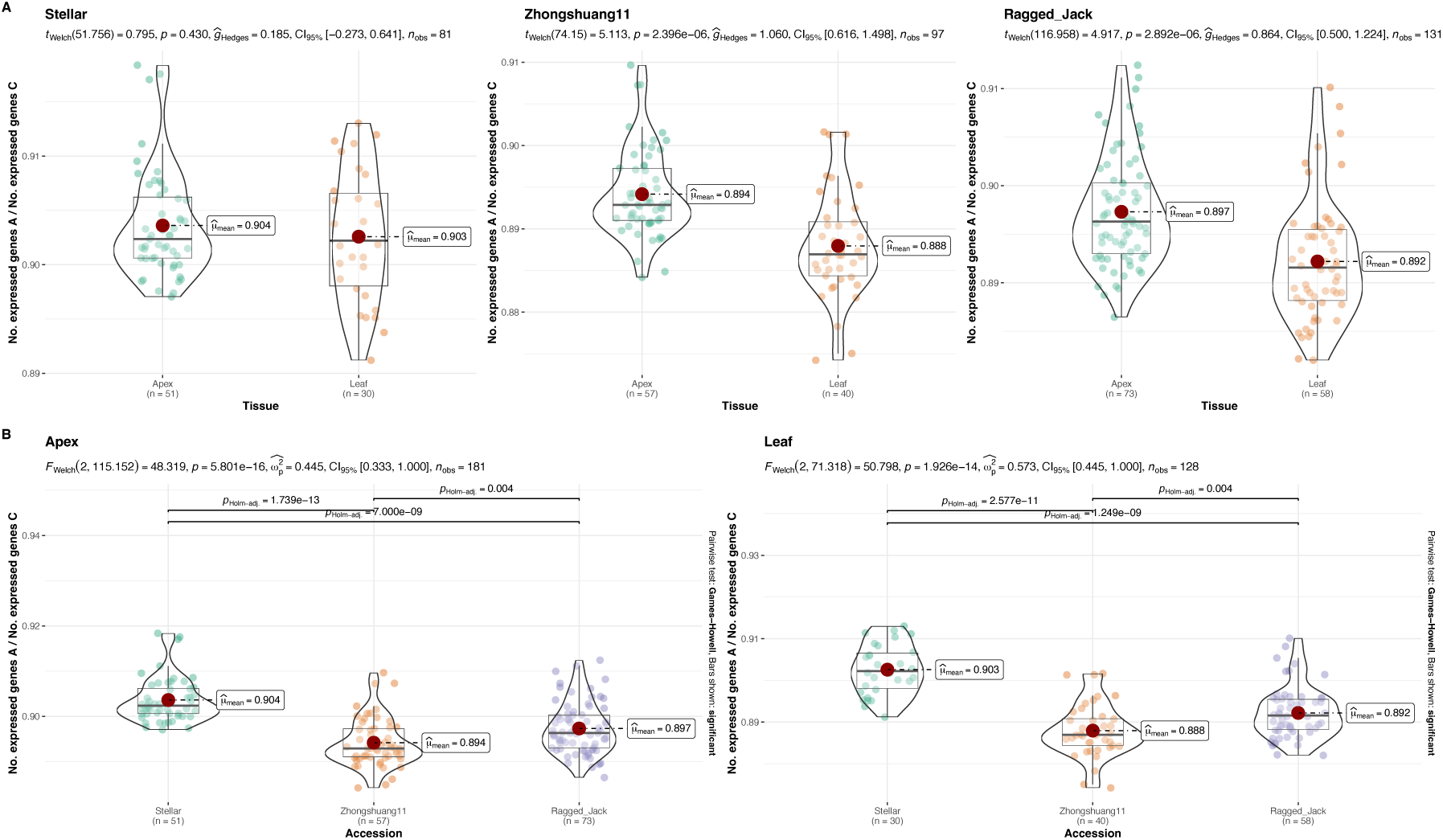
Comparison of the ratio of the number of expressed genes between tissues and cultivars. (A) Comparison between tissues for each of the cultivars, Ragged Jack, Zhongshuang 11, and Stellar. (B) Comparison between cultivars for each of the tissues.

**Figure S5.**
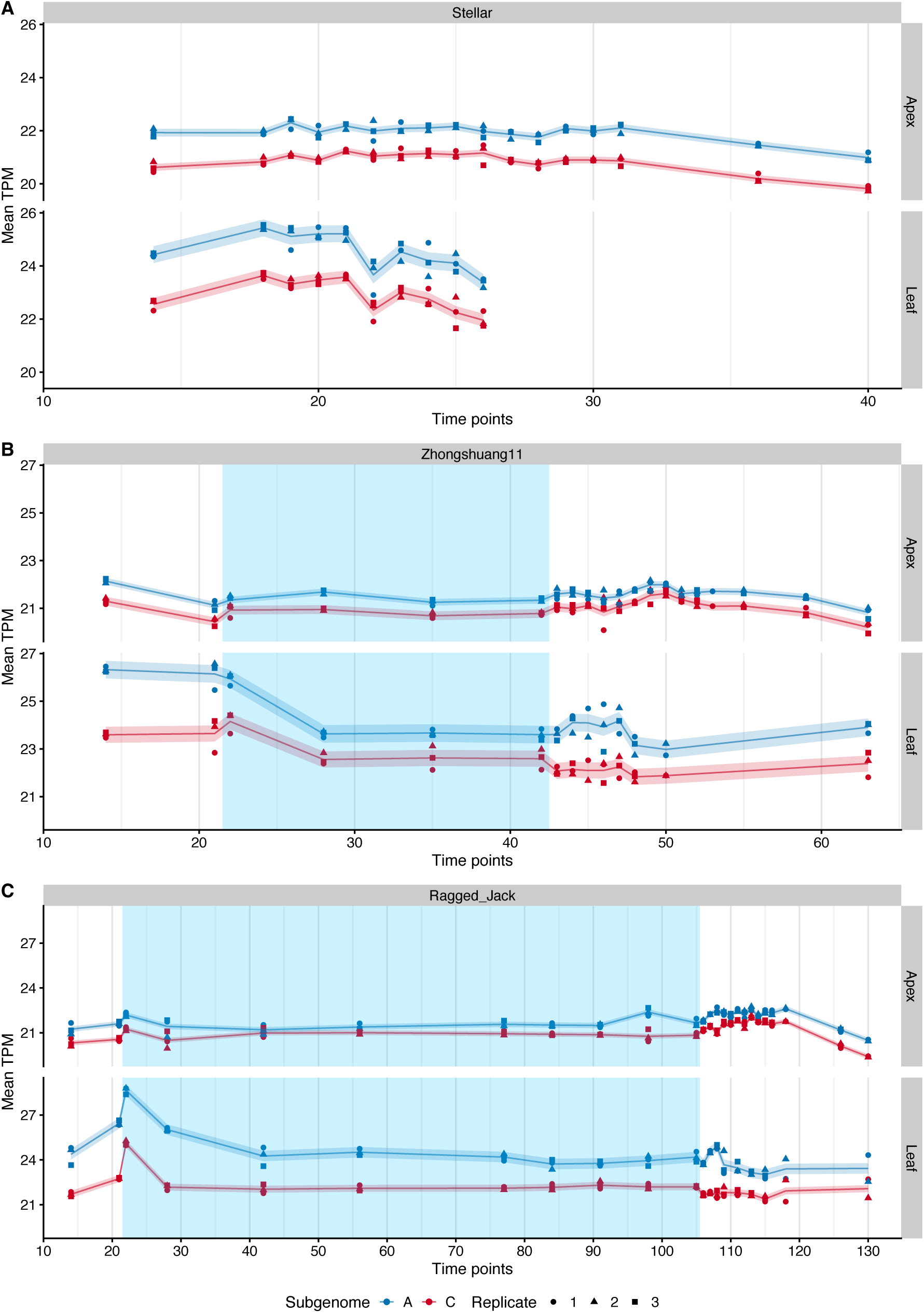
Mean expression of the genes shows that A subgenome dominates. The mean of the expression (TPM) for expressed genes by subgenome in both tissues, and in genotypes, Stellar (A), Zhongshuang11 (B), and Ragged Jack (C). The period of cold treatment is depicted by the blue shaded regions. A gene is considered to be expressed if its TPM is above 1. Each point represents a sample with the shape of the point indicating the replicate. The trend is shown with a thin line and the estimated standard deviation per time series by a shaded band (see Methods), with the colour indicating the subgenome (A: blue, C: red).

**Figure S6.**
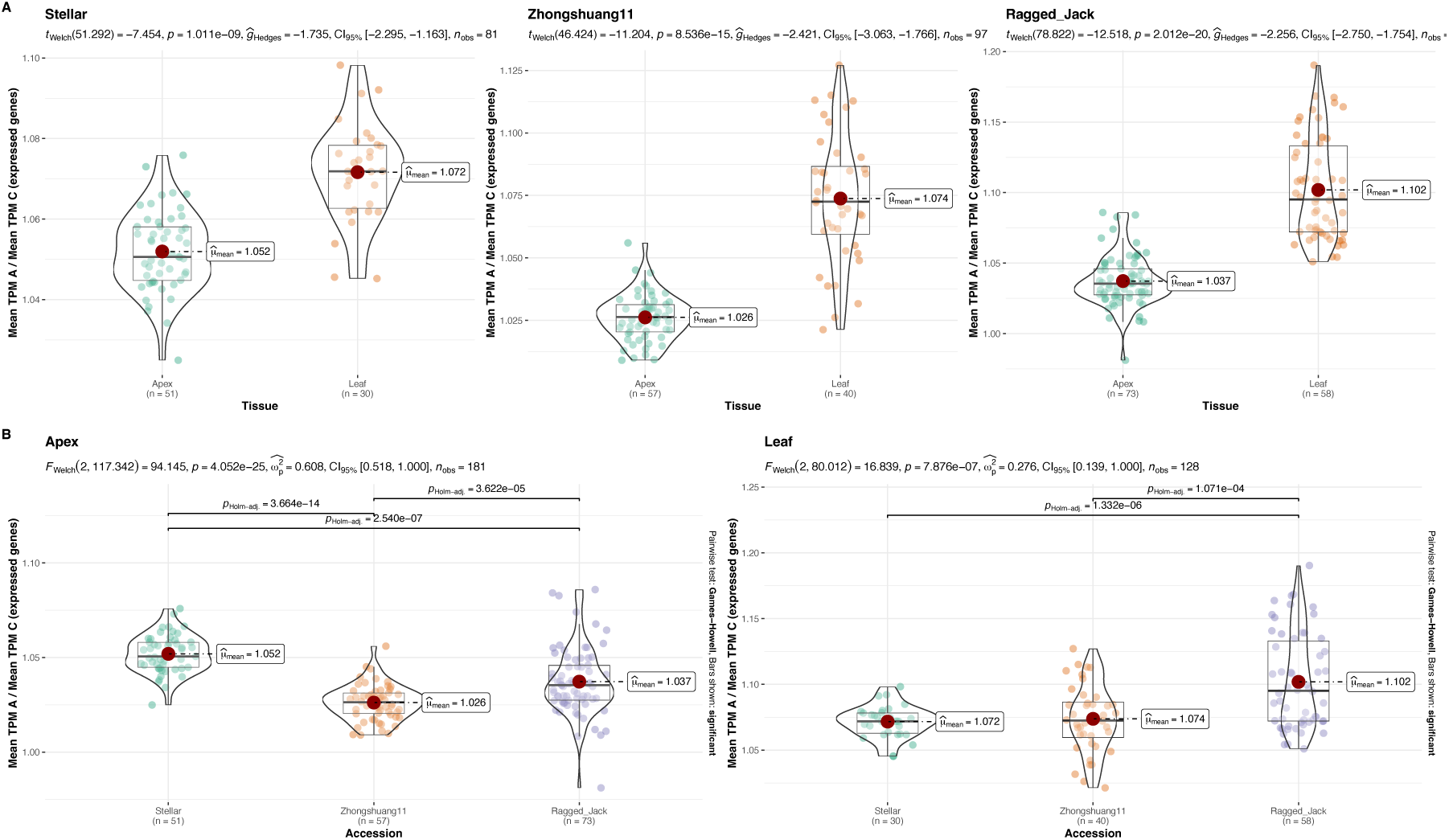
Comparison of the ratio of mean expression between tissues and cultivars. (A) Comparison between tissues for each of the cultivars, Ragged Jack, Zhongshuang 11, and Stellar. (B) Comparison between cultivars for each of the tissues.

**Figure S7.**
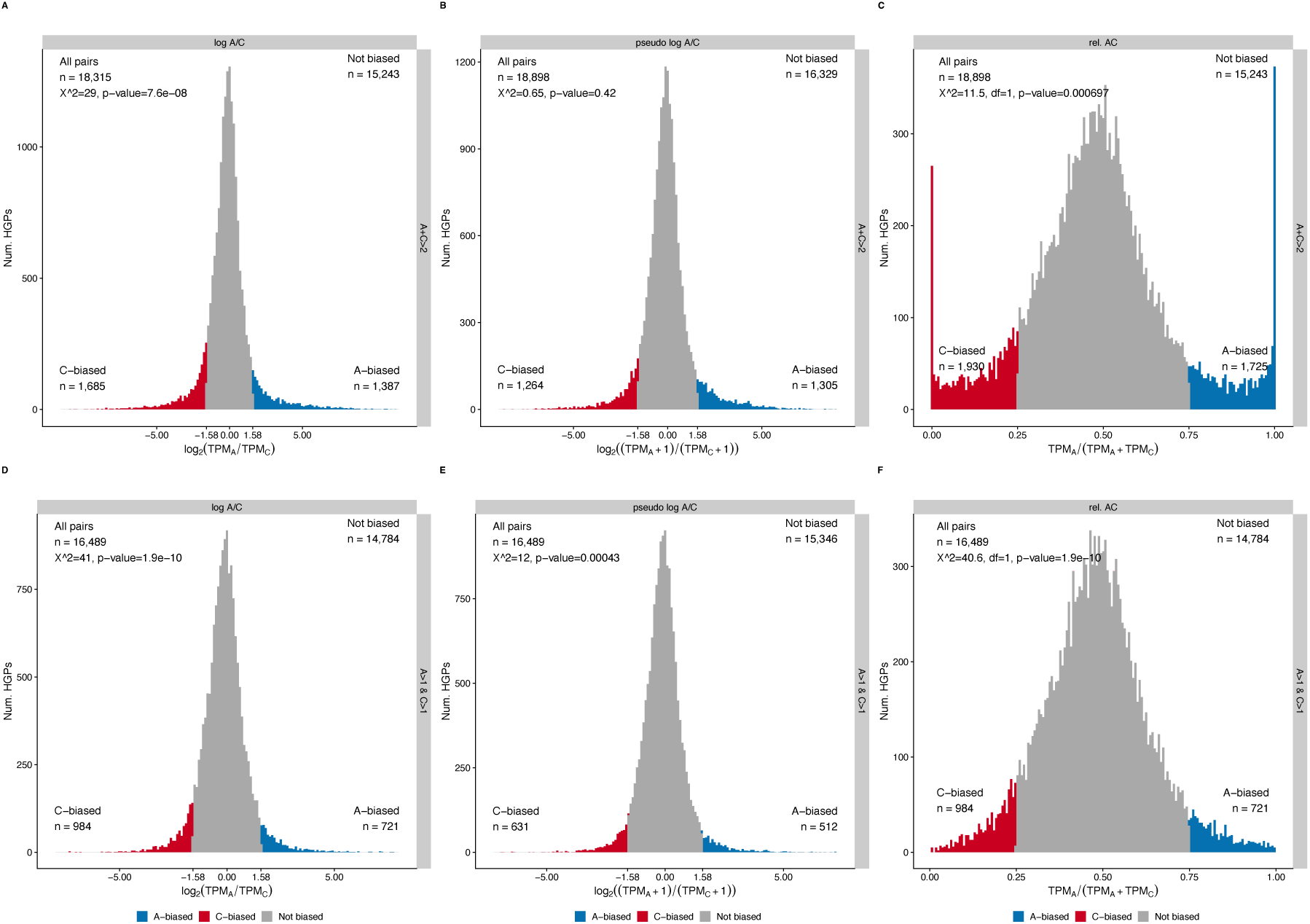
Comparison of subgenome bias classification methods to determine their impact on detecting bias. Distributions of the expression ratios for the homoeologue gene pairs (HGPs) of 14 day Stellar apex (replicate 1). The two expression filters are “A+C>2” (A, B and C) and “A>1 & C>1” (D, E and F). The three relative expression methods are “log A/C” (A, D), “pseudo log A/C” (B, E), and “rel. AC” (C, F). In each panel the total number of HGPs is given top-left with the number of HGPs that are considered to be not biased given top-right. The portion of the distribution that corresponds to the C-biased HGPs is to the left (red) with the number at the bottom-left, and the A-biased HGPs are to the right (blue) with the number shown at the bottom-right. The HGPs that fail to meet the bias criteria are considered to be unbiased (grey). The result of a 𝜒^!^-test on the numbers of A- and C-biased HGPs is shown at the top-left. The fold-change cut-off (FCC) is 3 throughout (log_2_(3) = 1.58 to 2 d.p.), which is equivalent to cut-offs of 1/4 and 3/4 for “rel. AC” (C, F). See Methods for full description of these expression filters and relative expression measures. The classification methods produce different numbers of biased HGPs in all cases except between (D) and (F). However, the methods producing the classification for (D) and (F) are equivalent for the expression filter and so the number of HGPs are expected to be the same. The number of C-biased HGPs is significantly higher than the number of A-biased HGPs in all cases except one (B) where not only are there more A-biased HGPs but the statistical test produces a not significant result.

**Figure S8.**
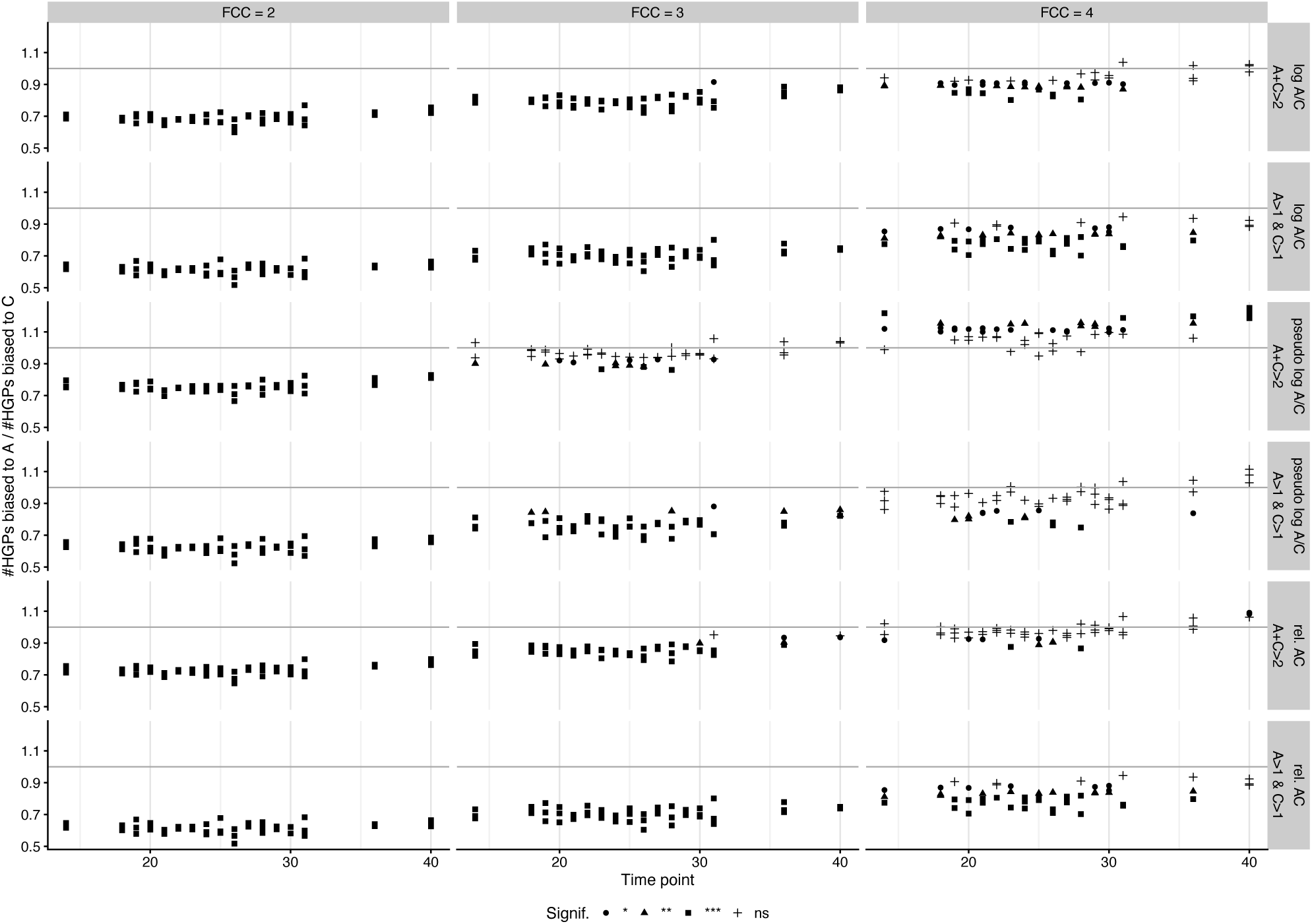
Assessing the effect of different methods of HGP bias classification across the apex time-course in Stellar. The ratio of the number of biased HGPs (A/C) (vertical axis) versus the time-point with each point representing a sample in the time-course, which was tested using the different methods. Bias was assigned to a HGP by a three step process: (1) filter the HGPs based on expression (“A+C>2” or “A>1 & C>1”), (2) calculate the relative expression (“log A/C”, “pseudo log A/C” or “rel. AC”), and finally, (3) label HGP as A-biased or C-based if the relative expression exceeds the fold-change cutoff (FCC) (see Methods). The numbers of HGPs that are labelled as biased to each subgenome were then tested ( 𝜒^!^ -test) with the significance indicated by the shape of the point (not significant (ns), <0.05 (*), <0.01 (**), <0.001 (***)). The ratio of the number of biased HGPs showed that, in most of the methods, more HGPs were biased to the C subgenome than to A. A different fold-change cut-off (FCC) could result in samples that were significantly biased to C for a low FCC either losing the significance for a higher FCC, or even reversing the result so that some samples were significantly biased to A. This reversal could also happen when we changed either the initial expression filter or the bias quantification method. This analysis highlights the possibility of different HGP bias classification methods producing different overall bias inferences.

**Figure S9.**
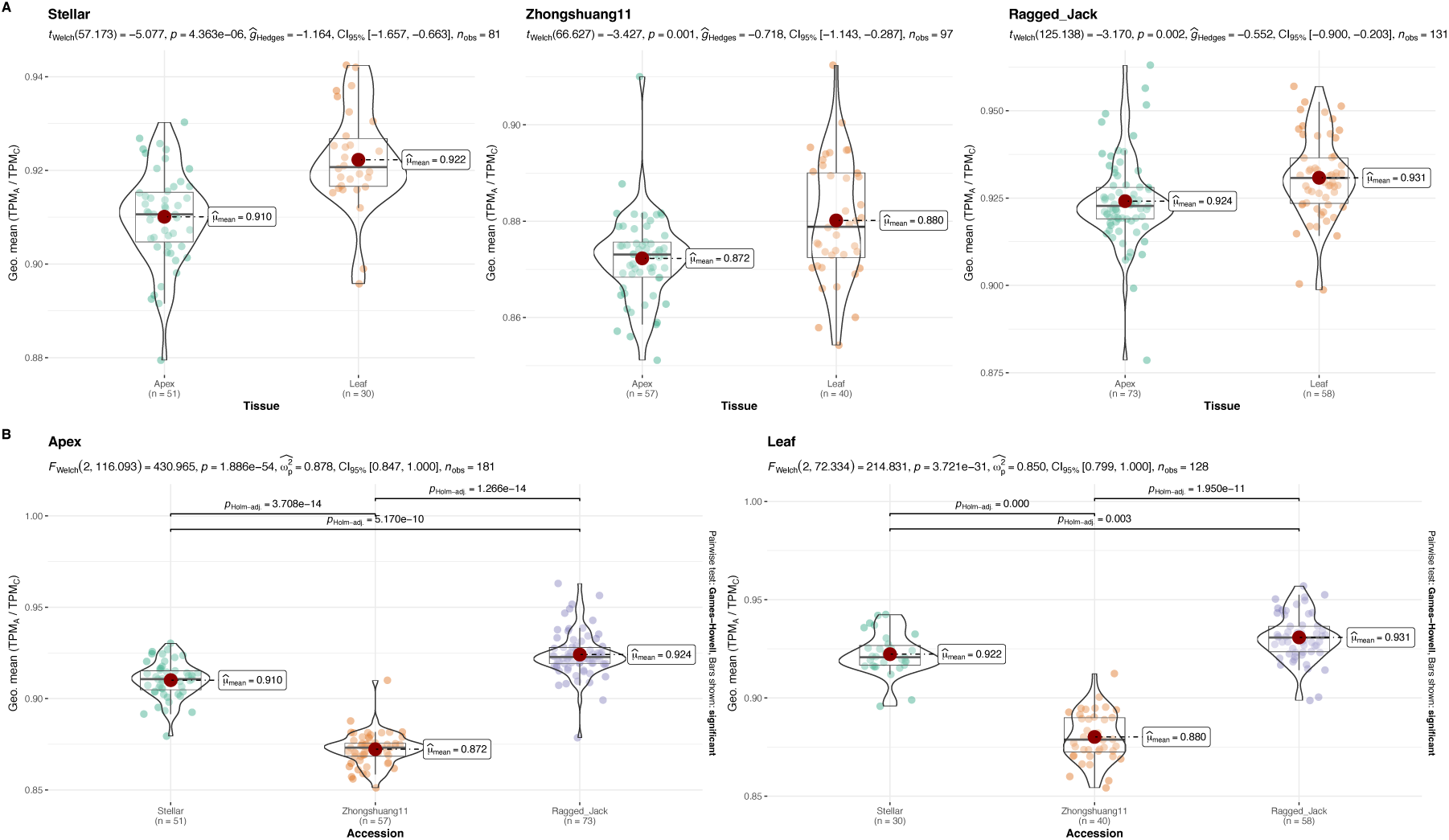
Comparison of mean expression ratios for the HGPs between tissues and cultivars. (A) Comparison between tissues for each of the cultivars, Ragged Jack, Zhongshuang 11, and Stellar. (B) Comparison between cultivars for each of the tissues. The mean biases for Stellar, Zhongshuang 11 and Ragged Jack were 9.9% C>A (7.5% – 13.7% C>A), 14.7% C>A (9.9% – 17.5% C>A) and 8.2% C>A (3.8% – 13.8% C>A) in the apex, respectively, and 8.4% C>A (6.1% – 11.6% C>A), 13.6% C>A (9.6% – 17.1% C>A), and 7.5% C>A (4.5% – 11.3% C>A) in the leaf, respectively.

**Figure S10.**
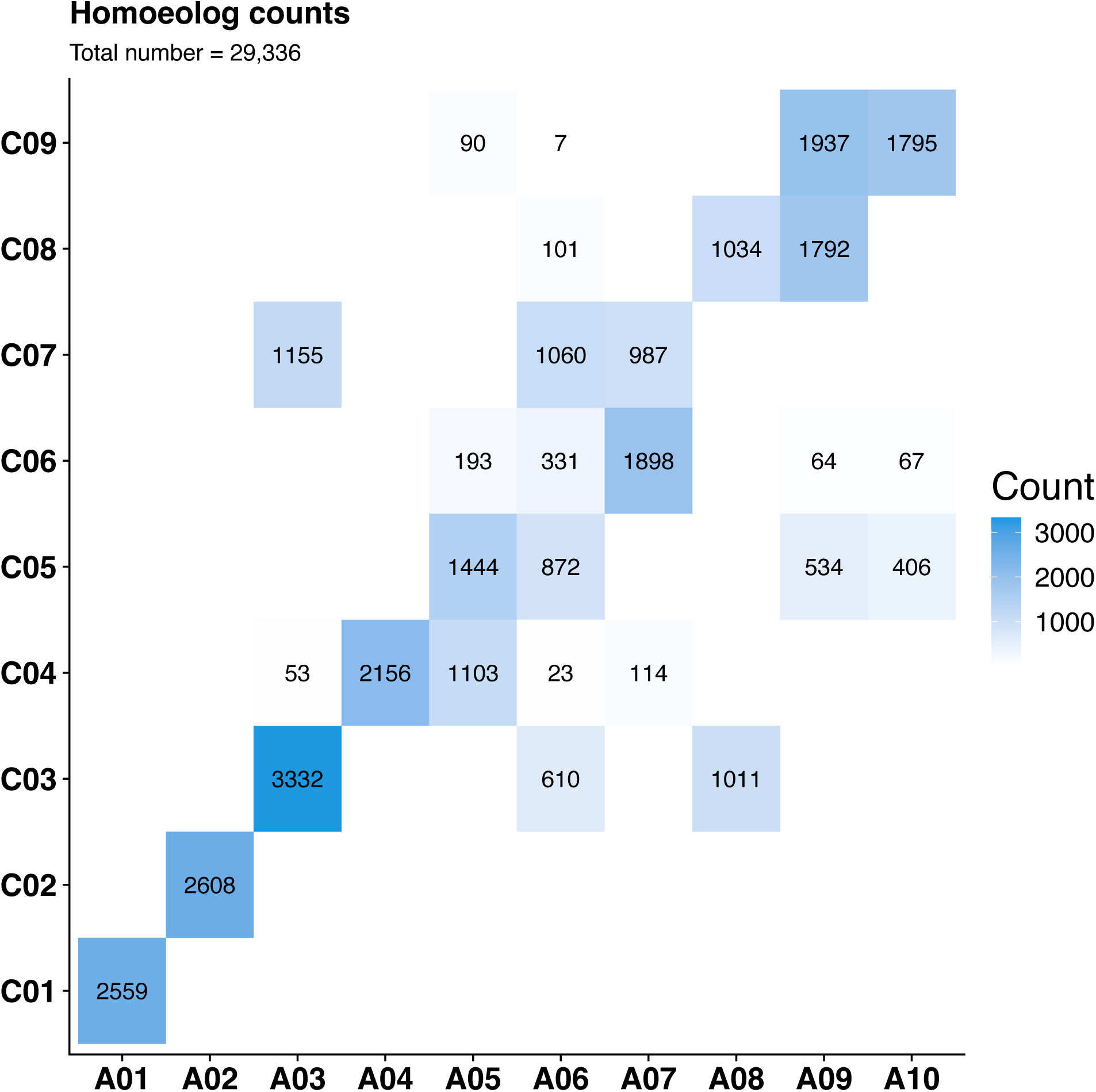
Number of homoeologue gene pairs (HGPs) between pairs of chromosomes on the two subgenomes. The results are from the “mmseqs2 easy-rbh” pipeline after taking synteny into account using GENESPACE (see Methods). The chromosomes for A and C are shown on the horizontal and vertical axes, respectively.

**Figure S11.**
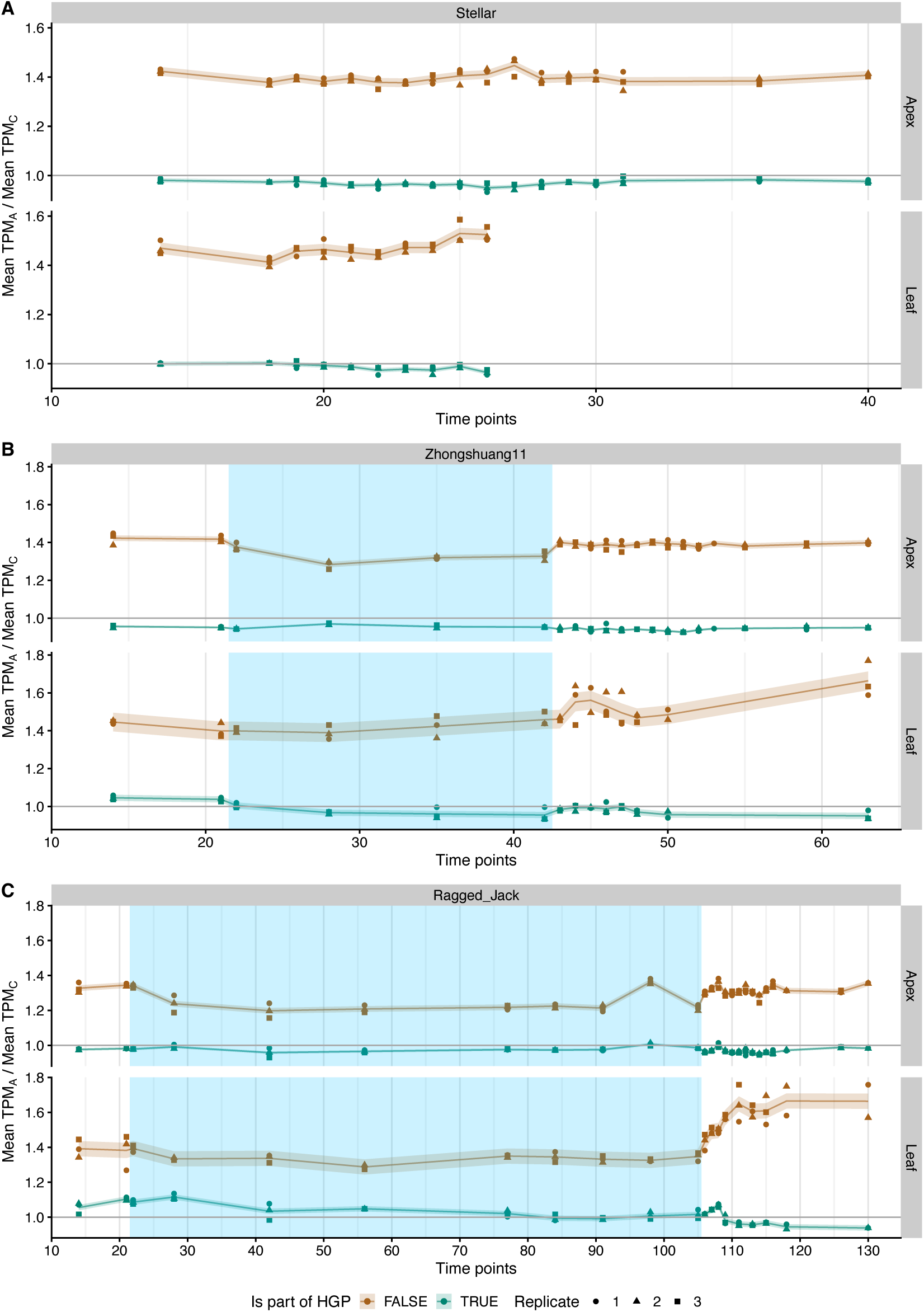
Mean expression ratio of the genes shows the difference in bias between genes that are part of homoeologue gene pair (HGP) and those that are not. The mean of the expression (TPM) for expressed genes by subgenome and whether the gene is part of a HGPin both tissues, and in genotypes, Stellar (A), Zhongshuang11 (B), and Ragged Jack (C). The period of cold treatment is depicted by the blue shaded regions. A gene is considered to be expressed if its TPM is above 1. Each point represents a sample with the shape of the point indicating the replicate. The trend is shown with a thin line and the estimated standard deviation per time series by a shaded band (see Methods), with the colour indicating whether the gene is a member of a HGP or not (Not part of HGP: brown, Part of HGP: teal).

